# Chronic restraint stress decreases resistance to punishment during cocaine self-administration via a dopamine D_1_-like receptor-mediated mechanism in prelimbic medial prefrontal cortex

**DOI:** 10.1101/2022.10.20.512776

**Authors:** Kevin T. Ball, Hunter Edson

**Affiliations:** Department of Psychology, Bloomsburg University of Pennsylvania, 400 E. 2^nd^ St., Bloomsburg, PA, 17815, USA

**Keywords:** CHRONIC STRESS, COCAINE, DOPAMINE, PRELIMBIC, PUNISHMENT-INDUCED ABSTINENCE, RELAPSE

## Abstract

We recently reported that male rats displayed greater resistance to punishment during cocaine self-administration compared to females. Moreover, daily restraint stress decreased this resistance in males, while having no effect in females. The purpose of the present study was to extend these findings by determining whether chronic stress-induced dopamine release in prelimbic medial prefrontal cortex mediates the effect of stress on punished cocaine self-administration. Thus, male rats were trained to press a lever for i.v. cocaine infusions (0.50 mg/kg/infusion) paired with a discrete tone + light cue in daily 3-hr sessions. Subsequently, 50% of the lever presses were punished by a mild footshock that gradually increased in intensity over 7 days. During the punishment phase, rats were exposed to a chronic restraint stress procedure (3 h/day) or control procedure (unstressed). Rats also received bilateral microinjections of the D_1_-like receptor antagonist SCH-23390 (0.25 μg/0.5 μl/side) or vehicle (0.5 μl/side) delivered to prelimbic cortex prior to daily treatments. Relapse tests were conducted 1 and 8 days after the last punishment session. Chronically stressed rats displayed reduced cocaine self-administration during punishment relative to unstressed rats, an effect prevented by co-administration of SCH-23390 to prelimbic cortex with daily restraint. Neither stress nor SCH-23390 treatment had significant effects on subsequent relapse-like behavior. These results establish a specific role for prelimbic D_1_-like receptors in chronic stress-induced suppression of punished cocaine self-administration in male rats. As such, these findings may inform novel methods to facilitate self-imposed abstinence in cocaine-dependent men.

## INTRODUCTION

In the treatment of addiction, abstinence is the goal. However, even when abstinence is achieved, vulnerability to relapse remains a major impediment to treatment success in the long term [1, 2]. Therefore, a great number of preclinical studies have been devoted to discovering the neural mechanisms driving both abstinence and relapse behaviors. Only a small number of preclinical studies, however, have assessed the influence of chronic stress on such behaviors, even though the clinical literature suggests there is a connection [3, 4]. Indeed, chronic stress induces long-lasting neurochemical and morphological alteration in medial prefrontal cortex (mPFC) [5, 6], a critical node in the circuitry driving many forms of abstinence and relapse in animal models [7-9].

We have investigated the effects of chronic stress on abstinence and relapse behavior by adding a chronic stress component to two rodent models of relapse: the classical reinstatement model [10, 11] and the punishment-induced voluntary abstinence model [12]. These two models differ in that the reinstatement model incorporates a form of “forced” abstinence whereby animals stop responding because the drug is no longer available (i.e., the response is extinguished), while the punishment model uses punishment of the drug self-administration response with contingent presentations of mild footshock to achieve “voluntary” abstinence. The latter procedure attempts to model the common situation in humans in which abstinence is self-imposed due to the negative consequences of continued drug use [13, 14]. This distinction is important because abstinence achieved using different methods may result in different outcomes as related to relapse behavior and underlying neural mechanisms [15, 16]. Indeed, we observed that chronic restraint stress had no effect on extinction of cocaine self-administration [17], but had a potentiating effect on abstinence achieved by punishment of the cocaine self-administration response [18].

The purpose of the present study was to build on these findings by assessing the involvement of dopamine D_1_-like receptors (D_1_Rs) in prelimbic (PL) mPFC in chronic stress’ ability to decrease resistance to punishment during cocaine self-administration. We used male rats because our previous results indicated that chronic stress had no effect on punished cocaine self-administration in female rats [18]. We focused on D_1_Rs in PL because 1) stress selectively activates the mesocortical dopamine pathway [19, 20], resulting in increases in extracellular dopamine in mPFC, 2) inactivation of PL (but not the more ventrally located infralimbic mPFC) blocks reinstatement of cocaine seeking [21-23], 3) dorsal mPFC injections of the D_1_R antagonist SCH-23390 attenuate reinstatement of cocaine seeking [24], 4) systemic co-administration of SCH-23390 with daily stress attenuates the potentiating effects of stress on future reinstatement of cocaine and palatable food seeking in male rats [17, 25-28], and 5) co-administration of SCH-23390 in PL, but not infralimbic, mPFC with daily stress attenuates stress’ ability to increase future cue-induced relapse to palatable food seeking in male rats [29].

## METHODS

### Subjects

Data were collected from male Sprague-Dawley rats (*n* = 24; Envigo) weighing 325-400 g at the commencement of procedures. Three rats were excluded from the study due to unreliable cocaine-reinforced responding during training (*n* = 2) and illness (*n* = 1). Rats were housed individually under standard laboratory conditions (12-hr light cycle from 7:00 AM to 7:00 PM) with *ad libitum* access to standard chow and water in their home cages and were weighed daily. All experiments were in compliance with NIH guidelines and were approved by the Bloomsburg University Institutional Animal Care and Use Committee.

### Apparatus

Testing occurred in standard modular operant conditioning chambers (Coulbourn Instruments, Whitehall, PA) that were housed in sound-attenuating, ventilated cubicles and connected to a PC with the Graphic State 4 software interface system (Coulbourn Instruments). Each chamber was equipped with an active and an inactive response lever. Chambers also included a house light, a row of multicolored LED cue lamps, a tone generator, and a fluid swivel with spring leash assembly connected to a counterbalanced arm assembly. A motor-driven syringe pump (Coulbourn Instruments) for drug delivery was located outside of each cubicle.

### Surgery

Rats underwent surgery to implant a jugular catheter. Animals were anesthetized with ketamine HCl (90 mg/kg; i.m.) and xylazine HCl (10 mg/kg; i.m.), with supplemental injections as needed. A catheter, made of silastic tubing (0.51 m I.D. x 0.94 mm O.D.; Dow Corning, Midland, USA) attached to a modified 22-gauge cannula-guide connector assembly (Plastics One, Roanoke, VA), was inserted into the right jugular vein. The catheter was routed subcutaneously and exited at the scapula of the skull. Following catheterization, rats were fixed in a stereotaxic apparatus and two guide cannulas (22 gauge; Plastics One) were bilaterally implanted 1.0 mm above the PL (coordinates: AP: +3.2, ML: ±1.3, and DV –3.0 relative to bregma, midline, and skull surface, respectively, with a 10° angle away from midline). Coordinates were based on our previous research [29, 30]. Five stainless steel screws were implanted in the skull for support, and the cannulas and catheter were held in place with dental cement. A dummy cannula was inserted into each guide cannula to prevent blockage. Catheters were flushed with 0.1 ml (10 mg/ml; i.v.) of gentamycin (Lonza, Walkersville, MD) and heparinized physiological saline (30 U/ml heparin) once daily. Injections of 0.1 ml Brevital (1%) were used to test catheter patency if rats did not acquire cocaine self-administration after 7 days or if cocaine infusions earned decreased by > 20% from the previous day. Loss of muscle tone within 5 s after injection indicates a patent catheter.

### Training phase

Following 1 week of recovery from surgery, rats were trained to press the lever for cocaine HCl (NIDA Drug Supply Program, Research Triangle Park, NC) in daily 3-hr sessions. During these sessions rats responded on a fixed ratio-1 (FR-1) schedule under which responses on the active lever resulted in an intravenous infusion of 0.50 mg/kg of cocaine (weight of salt; dissolved in 0.9% sterile saline) accompanied by conditioned stimuli (CS), which consisted of a tone + flashing cue light compound stimulus presented for 5 s. Drug concentration (5.0 mg/ml) and pump delivery rate (10 µl/s) were kept constant, and dose of drug (0.50 mg/kg/infusion) was controlled by varying the duration of pump action (i.e., volume of injected solution) based on body weight. Delivery of the drug and CS was followed by a 20 s time-out period signaled by illumination of the house light. During cocaine infusions and time-outs responses were recorded but had no programmed consequences. The criterion of ≥ 12 cocaine infusions in one session was used to indicate acquisition. After the acquisition criterion was reached, rats self-administered cocaine for 9 more daily sessions. See Fig. 1a for a timeline of the experiment.

**Fig. 1.**
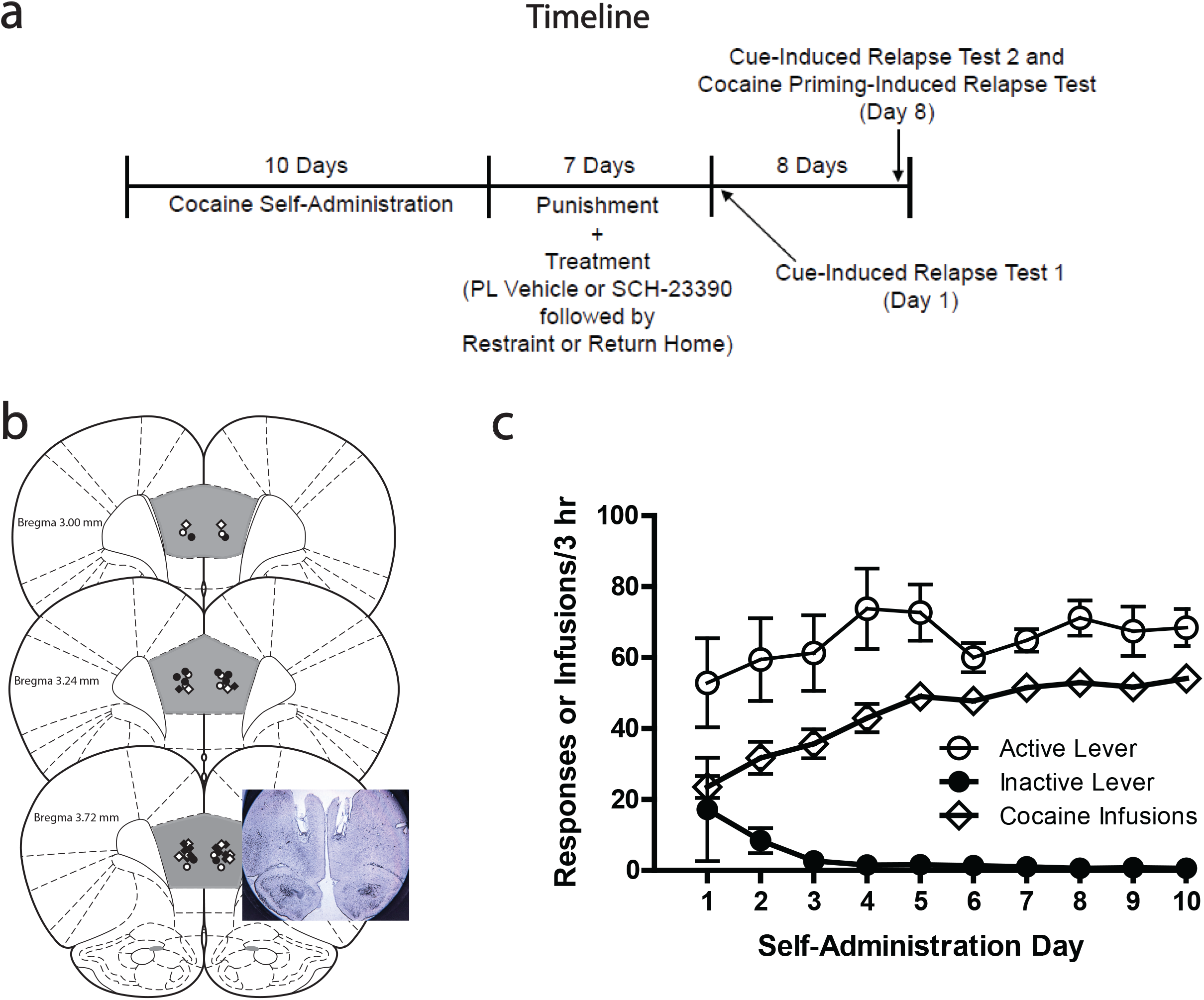
(a) Timeline of the experiment. (b) Schematic illustration (modified from Paxinos and Watson, 2005 [46]) of coronal sections from the rat brain showing approximate injection sites in PL. Open and filled circles represent the injection sites in VEH + unstressed and VEH + stressed treatment groups, respectively, and open and filled diamonds represent the injection sites in SCH-23390 + unstressed and SCH-23390 + stressed treatment groups, respectively. *n* = 5-6/treatment group. *Inset*: Photomicrograph of coronal section illustrating cannula tracts from PL placement. Notch indicates right hemisphere. (c) Acquisition of cocaine self-administration. Active lever presses were reinforced on a FR-1 20-s timeout schedule in daily 3-h sessions. Delivery of cocaine (0.50 mg/kg/infusion) was accompanied by a tone + flashing cue light CS presented for 5 s. Day 1 represents responding on the first day that the acquisition criterion was met. Data represent mean ± SEM.

### Punishment phase

On the day following the last self-administration session, rats continued cocaine self-administration under the same FR-1 20-s timeout schedule, except that 50% of the reinforced lever presses resulted in the concurrent delivery of a 0.5-s footshock through the grid floor. Footshock began at 0.12 mA and increased by 0.06 mA every day to a final value of 0.48 mA (a total of 7 punishment sessions). These parameters are based on previous studies of punishment-induced abstinence from drug seeking [12, 18, 31]. Punished responses continued to produce the CS and 0.50 mg/kg cocaine.

### Chronic restraint stress and intracranial injections

Pairs of rats were matched based on mean cocaine self-administration across the last 3 days of the training phase. One rat from each pair was randomly assigned to the stress condition. Beginning on the first day of the punishment phase (∼30 min after punishment sessions) and continuing until the last day, stressed rats (*n* = 11) were exposed to daily restraint stress for 3 h by placement in plastic semi-cylindrical restrainers (8.57 cm diameter x 21.59 cm length; Braintree Scientific). Restraint stress is a well-established procedure that produces significant increases in plasma corticosterone in rats [32, 33]. Unstressed rats (*n* = 10) were handled for 5 min each day and then returned to their home cage. Immediately prior to exposure to restraint stress or the control condition, rats received bilateral microinjections of SCH-23390 (0.25 μg/0.5 μl/side) or saline (0.5 μl/side) delivered to PL mPFC. This resulted in four treatment groups: vehicle (VEH) + unstressed (*n* = 5), VEH + stressed (*n* = 5), SCH-23390 + unstressed (*n* = 5), and SCH-23390 + stressed (*n* = 6). PL infusions of this volume and concentration of SCH-23390 were chosen because previous research showed that it prevented both PL pyramidal cell dendritic retraction [34] and future cue-induced relapse to palatable food seeking [29] in rats undergoing daily restraint stress. Microinjections were delivered through 28-gauge internal cannulas that extended 1 mm below the guide cannula by an infusion pump with 10 μl syringes over a 1-min period. Syringes were connected to internal cannulas with polyethylene tubing. Internal cannulas were left in place for 1 min after injections to allow for drug diffusion.

### Relapse tests

Relapse tests in the presence of the cocaine-associated cues were conducted on the 1^st^ (Test 1) and 8^th^ (Test 2) days after the last session of the punishment phase. Active lever presses resulted in the delivery of the CS that was previously paired with cocaine delivery on an FR-1 20-s timeout schedule, but not cocaine. Test 1 was a single 1-h session. Test 2 consisted of five 1-h sessions (separated by 5 min) that were each identical to Test 1 except that rats were given 0.0, 5.0, and 10.0 mg/kg cocaine injections (i.p.) immediately before the start of sessions 3, 4, and 5, respectively. Doses of cocaine were given in an ascending order to minimize carry-over effects of residual cocaine. This within-session procedure is based on previous studies with cocaine priming [35, 36]. Between test days, rats remained in their home cages and were handled every other day. Relapse was assessed as follows: Test 1 (1 h) and session 1 of Test 2 (1 h) were used to assess cue-induced relapse, as well as incubation of craving [37]. Sessions 3, 4, and 5 of Test 2 were used to assess cocaine priming-induced relapse. Data from session 2 were not analyzed; this session served as an additional period of time for lever pressing to diminish prior to session 3 (cocaine priming control).

### Histology

Upon completion of experiments, rats were deeply anesthetized with a commercial euthanasia solution containing pentobarbital sodium (390 mg/ml) + phenytoin sodium (50 mg/ml) and then perfused transcardially with 0.9% saline followed by 10% formaline. Brains were removed and stored in formalin solution (∼1 week) followed by 30% sucrose/10% formalin solution (∼1 week), then frozen, cut (60 μm sections), mounted on slides, and stained with cresylecht violet to verify microinjection sites. An observer blind to treatment conditions determined cannula placements. Fig. 1b shows the positions of microinjection sites.

### Statistical analyses

Data were analyzed using mixed factorial ANOVAs. The main dependent variables were number of cocaine infusions (during self-administration and punishment phases) and number of lever responses (during relapse testing). Change in body weight across chronic treatment days also was used as a dependent variable in order to verify the physiological relevance of the stress manipulation. The between-subjects factors were SCH-23390 dose (0.0 or 0.25 μg/side) and stress condition (stressed or unstressed). The within-subjects factors were self-administration day (1-10 and 0-7 for training and punishment phases, respectively), chronic treatment day (2-7 for body weight data), lever (active and inactive), abstinence day [1 (Test 1) and 8 (Test 2) for cue-induced relapse], and cocaine priming dose (0.0, 5.0, and 10.0 mg/kg for cocaine priming-induced relapse). Because the factorial ANOVAs resulted in multiple main and interaction effects, we report only significant *F* values that are important for interpretation.

## RESULTS

### Training phase

As shown in Fig. 1c, number of cocaine infusions increased over days [main effect of day, *F*(9, 180) = 23.35, *p* < .001] following acquisition (Day 1). Mean (±SEM) number of infusions on Day 10 were not significantly different among rats assigned to different treatment groups [55.40 (2.82), 52.00 (4.91), 54.80 (3.18), and 54.17 (2.41) for VEH + unstressed, VEH + stressed, SCH-23390 + unstressed, and SCH-23390 + stressed groups, respectively].

### Punishment Phase

#### Body weight

As expected, restraint was a stressor, as indicated by body weight loss [main effect of stress, *F*(1, 17) = 43.62, *p* < .001; day X stress interaction, *F*(5, 85) = 11.87, *p* < .001; see Fig. 2a]. PL SCH-23390 injections had no effect on body weight in either stress condition.

**Fig. 2.**
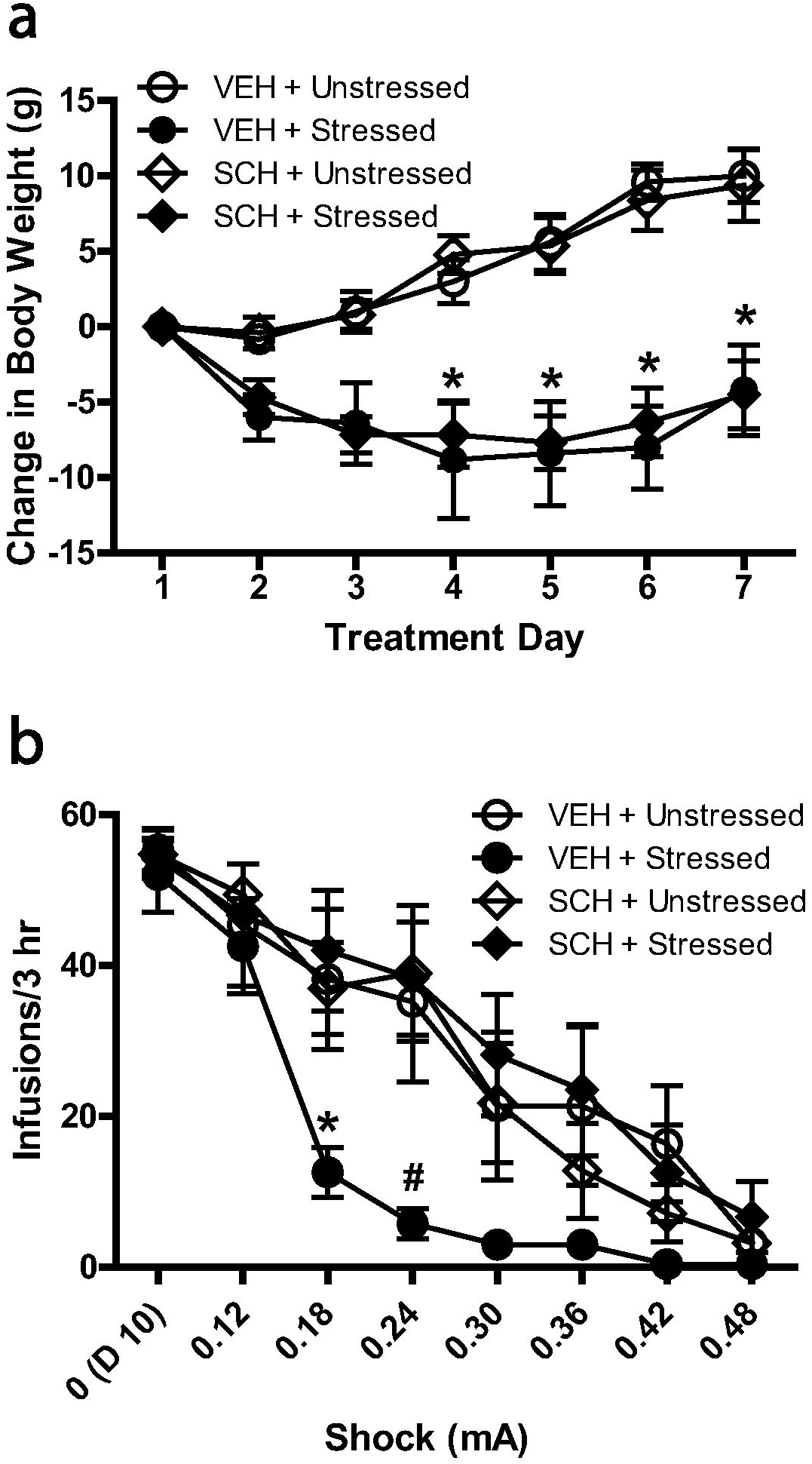
(a) Mean (± SEM) change in body weight in chronically stressed and unstressed rats receiving saline (0.5 μl/side) or SCH-23390 (0.25 μg/0.5 μl/side) infusions targeting PL mPFC. Each day following punishment sessions, rats were given their assigned PL injection and placed in restrainers (3 h/day) or returned to their home cage. \**p* < .01 compared with unstressed groups, Bonferroni post-test. (b) Mean (± SEM) cocaine infusions earned during the punishment phase. During this phase, shock intensity was gradually increased from 0.12 to 0.48 mA over 7 days, and 50% of reinforced lever presses were accompanied by concurrent footshock. \**p* < .05 compared with SCH + stressed group and **#***p* < .05 compared with all other treatment groups, Bonferroni post-test.

#### Cocaine self-administration

As shown in Fig. 2b, number of cocaine infusions decreased over days during the punishment phase for all treatment groups [main effect of day, *F*(7, 119) = 65.77, *p* < .001]; however, the VEH + stressed group was more sensitive to punishment compared to all other groups as indicated by reduced cocaine self-administration beginning on punishment Day 2, (0.18 mA of shock; the day after the first stress exposure). By contrast, stressed and unstressed rats receiving PL SCH-23390 displayed cocaine intake similar to the VEH + unstressed group [day X SCH-23390 X stress interaction, *F*(7, 119) = 2.13, *p* < .05]. By punishment Day 7 (0.48 mA of shock), all treatment groups displayed similarly low levels of cocaine intake.

#### Relapse

As shown in Fig. 3a, responding for cocaine-associated cues was not significantly different among treatment groups on abstinence Day 1 or 8. There also was no significant difference in responding overall from Day 1 to 8. As shown in Fig. 3b, although active, but not inactive, lever pressing increased as a function of cocaine priming dose [main effect of priming dose, *F*(2, 34) = 6.76, *p* < .01; lever X priming dose interaction, *F*(2, 34) = 7.58, *p* < .01], there were no effects of prior PL SCH-23390 or stress on this form of relapse.

**Fig. 3.**
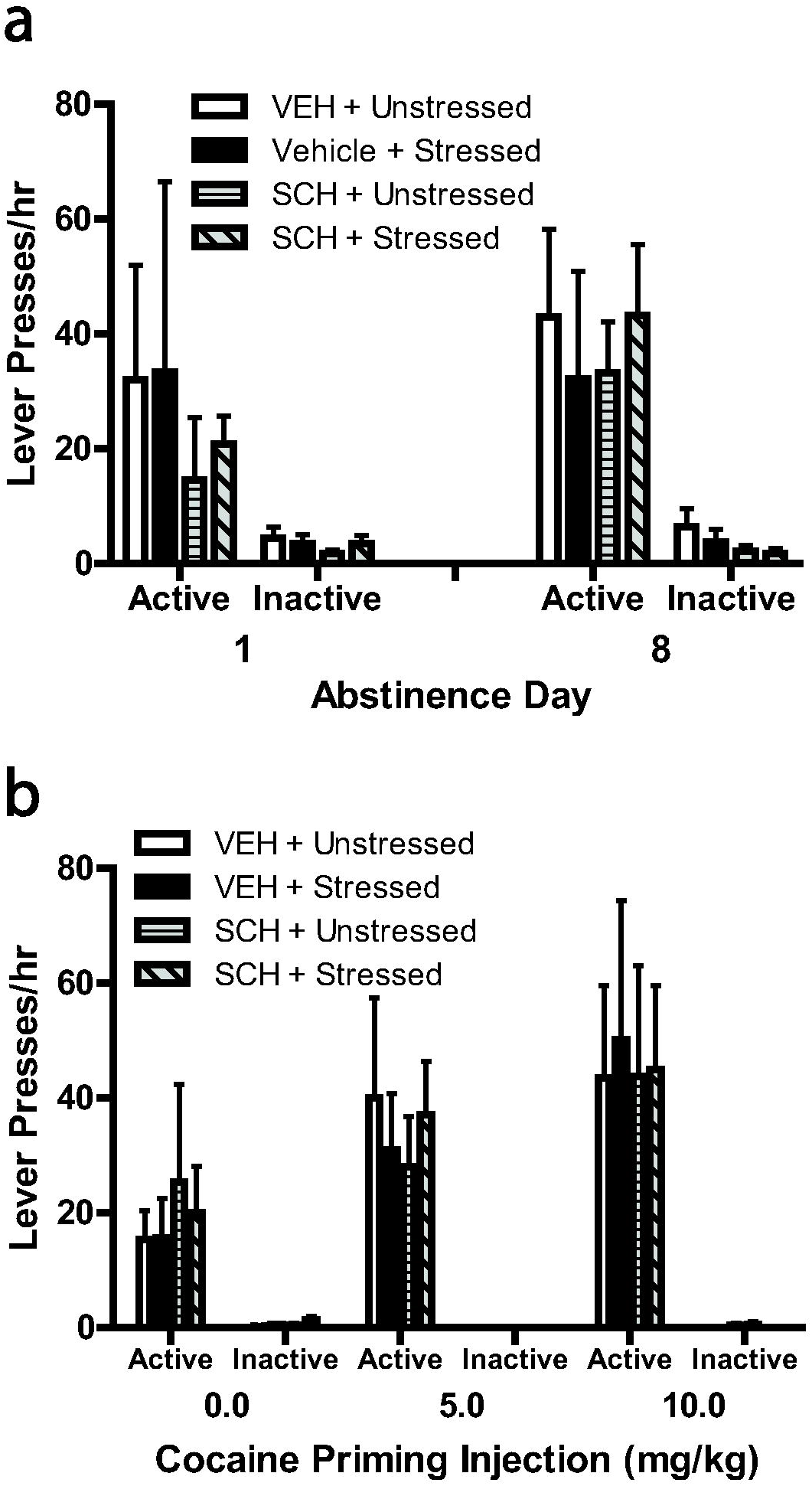
(a) Mean (+ SEM) active and inactive lever presses during cue-induced relapse tests. Contingencies during these tests were identical to self-administration sessions except that the primary reinforcer (cocaine) was not delivered. Test 1 occurred on the day following the last punishment session. Test 2 occurred 7 days later. (b) Mean (+ SEM) active and inactive lever presses during cocaine priming-induced relapse tests. The testing consisted of three consecutive 1-h sessions that were identical to self-administration sessions, except that lever presses did not lead to cocaine delivery, and rats were given 0.0, 5.0, and 10.0 mg/kg cocaine injections (i.p.) at the beginning of hours 1, 2, and 3, respectively.

## DISCUSSION

We tested the effect of daily restraint stress on punished cocaine self-administration and subsequent cue-and cocaine priming-induced relapse in male rats. Supporting our previous results [18], we found that stress caused a decrease in resistance to punishment, while having no effect on later cue-or cocaine priming-induced relapse. We extend these findings by showing that co-administration of SCH-23390 delivered to PL cortex with daily restraint prevented the effect of stress on punished cocaine self-administration. Thus, these results establish a specific role for PL D_1_Rs in chronic stress-induced suppression of punished cocaine self-administration.

It is noteworthy that, as we reported previously [18], the effects of restraint on punished cocaine self-administration were observed after just one restraint session (i.e., on Day 2 of punishment). Because the first restraint session ended ∼18 hr before the second punishment session, the effects of this acute stress-induced PL D_1_R stimulation must have resulted in some form of enduring neuroplasticity. Although any discussion of mechanisms downstream from D_1_R stimulation is speculative, there is ample evidence of dopaminergic modulation of long-term synaptic plasticity in mPFC involving D_1_Rs [38]. There also is evidence that one 40-min session of intermittent footshock stress results in PL pyramidal cell dendritic retraction 24 h later [39]. Although we are not aware of any studies that have assessed the role of dopamine in acute stress-induced dendritic remodeling in PL, such changes induced by chronic restraint stress are prevented by co-administration of PL SCH-23390 with the stress [34]. Given that exposure to cocaine itself, however, causes lasting neurochemical, structural, and functional changes in mPFC [40-42], the specific stress-induced mPFC plasticity that occurs in animals with a history of cocaine self-administration, as well as the role of D_1_Rs in such plasticity, awaits further investigation.

The ability of PL SCH-23390 to prevent stress-induced decreases in punished cocaine self-administration does not appear to be due to a reduction in the hypothalamic-pituitary-adrenal axis response to stress for two reasons. First, we observed similar weight loss in the SCH-23390 + stressed and VEH + stressed groups. Weight loss occurs following exposure to a variety of stressors and is commonly used as a verification of the stress manipulation in preclinical studies [43]. Second, Lin et al. reported that PL D_1_R blockade with SCH-23390 had no effect on acute restraint stress-induced increases in plasma corticosterone levels [34]. Thus, the observed effects appear to be due to the ability of SCH-23390 to block plasticity in PL cortex due to stress-induced dopamine release in this region, rather than a reduction in the overall “stressfulness” of the manipulation. Given that D_1_Rs in mPFC mediate dopamine effects on cognitive functioning [44], and also that the administration of SCH-23390 following daily extinction training results in impaired extinction of cocaine seeking [17] and cocaine conditioned place preference [45], it is possible that stress-induced dopamine release in PL cortex helped to consolidate response-outcome associations to facilitate punishment-induced abstinence from cocaine self-administration, and that PL D_1_R blockade inhibited this process.

A concern with regard to the interpretation of our results is that SCH-23390 may have diffused away from PL to neighboring regions of mPFC. Although we cannot completely rule out this possibility, it seems very unlikely given our previous results using targeted, daily injections of SCH-23390. Thus, using the same SCH-23390 dose, injection volume, and daily treatment schedule as in the present study, PL injections combined with daily restraint stress prevented stress-induced increases in subsequent relapse to palatable food seeking, whereas injections targeting neighboring infralimbic cortex were without effect [29]. Such findings are consistent with localized effects of SCH-23390 within mPFC regions.

In sum, we report a critical role for PL D_1_Rs in the ability of stress to suppress punished cocaine self-administration in male rats, extending our previous findings that D_1_R antagonism prevents chronic stress’ ability to potentiate future relapse-like behavior following forced abstinence from cocaine and palatable food seeking in male rats [17, 25-29]. Collectively, these results provide evidence that chronic stress can both increase and decrease reward seeking, but that D_1_R activation is required in both cases. Thus, PL dopamine should be an important focus of future mechanistic studies on the relationships between chronic stress, abstinence, and relapse.

## Acknowledgements

This work was supported by the National Institutes of Health (NIDA R15 DA035432 to KTB). Cocaine was generously provided by the National Institute on Drug Abuse.

